# Systemic analysis of putative SARS-CoV-2 entry and processing genes in cardiovascular tissues identifies a positive correlation of BSG with age in endothelial cells

**DOI:** 10.1101/2020.06.23.165324

**Authors:** Blerina Ahmetaj-Shala, Ricky Vaja, Santosh S Atanur, Peter M. George, Nicholas S. Kirkby, Jane A. Mitchell

**Affiliations:** Cardiorespiratory Interface, National Heart and Lung Institute, Imperial College London, SW7 2AZ; Department of Metabolism, Digestion and Reproduction, Faculty of Medicine, Imperial College London; Institute of Translational Medicine and Therapeutics (ITMAT) Data Science Group, NIHR, BRC, Imperial College London; Interstitial Lung Disease Unit, National Heart and Lung Institute, Imperial College London, Royal Brompton and Harefield NHS Foundation Trust, Sydney Street, London, SW3 6NP, UK

## Abstract

COVID-19, caused by severe acute respiratory syndrome coronavirus-2 (SARS-CoV-2) has rapidly spread throughout the world with unprecedented global healthcare and socio-economic consequences. There is now an established secondary syndrome of COVID-19 characterised by thrombosis, vascular dysfunction and hypertension, seen in those most severely affected. Advancing age in adults is the single most significant risk factor for hospitalisation and death with COVID-19. In light of the cardiovascular/thrombotic sequalae associated with severe COVID-19 disease and the overwhelming risk that increased age carries, in this study, our aim was to obtain mechanistic insight by interrogating gene expression profiles in cardiovascular tissues and cells. Our focus was on the two putative receptors for SARS-CoV-2, *ACE2* and *BSG* along with a selected range of genes thought to be involved in virus binding/processing. In this study we have made four important observations: (i)Cardiovascular tissues and/or endothelial cells express the required genes for SARS-CoV-2 infection, (ii) SASR-CoV-2 receptor pathways, *ACE2*/*TMPRSS2* and *BSG*/*PPIB(A*) polarise to lung/epithelium and vessel/endothelium respectively, (iii) expression of SARS-CoV-2 host genes are, on the whole, relatively stable with age and (iv) notable exceptions were *ACE2* which decreases with age in some tissues and *BSG* which increases with age in endothelial cells. Our data support the idea that that BSG is the dominate pathway utilised by SARS-CoV-2 in endothelial cells and are the first to demonstrate a positive correlation with age. We suggest BSG expression in the vasculature is a critical driver which explains the heightened risk of severe disease and death observed in those >40 years of age. Since BSG is utilised by other pathogens our findings have implications beyond the current pandemic. Finally, because BSG is functions in a range of cardiovascular diseases and fibrosis, our observations may have relevance to our understanding of the diseases associated with aging.

## Introduction

COVID-19, caused by severe acute respiratory syndrome coronavirus-2 (SARS-CoV-2), was first reported in Wuhan, China in December 2019. SARS-CoV-2 is relatively contagious and whilst producing mild symptoms in the majority of people, can progress to severe or fatal disease in susceptible individuals. As such the virus has rapidly spread throughout the world with unprecedented global healthcare and socio-economic consequences.

SARS-CoV-2 is related to SARS-CoV and Middle East respiratory syndrome coronavirus (MERS-CoV) which caused respiratory epidemics in 2003 and 2012 respectively. Based on what was known about human host interactions with SARS-CoV and MERS-CoV along with recent research using SARS-CoV-2 tools, a list of key entry and processing genes utilised by the virus to infect host cells has been defined. SARS-CoV-2 enters host cells by binding of the spike protein with two putative receptors; ACE2^1^ and BSG (also known as Basigin, CD147 or EMMPRIN)^2,3^. For viral entry by ACE2, it is thought that the SARS-CoV-2 spike protein is primed and ACE2 cleaved, by the cellular serine proteases TMPRSS2^1^ and ADAM17. FURIN cleaves viral enveloping proteins providing another putative priming step for the spike protein of SARS-COV-2^4^. For viral entry via BSG, less is known regarding specific receptor/viral processing partners for SARS-CoV-2. However, for SARS-CoV^5^, HIV^6^ and the measles virus^7^, respectively, peptidylprolyl isomerase A (PPIA; also knowns as cyclophilin A) and peptidylprolyl isomerase B (PPIB; also known as cyclophilin B), which are natural ligands for BSG, incorporate into virus and facilitate binding to BSG. Similarly, PPIB forms a complex with the malaria pathogen (*Plasmodium falciparum merozoites*) and BSG to facilitate infection of red blood cells^8^. Intracellular processing of SARS-CoV-2 spike protein is thought to involve the lysosomal cysteine proteases cathepsin B/L (CTSL, CTSB) which, can also substitute for TMPRSS2 in some cells^1^.

Initial infection with SARS-CoV-2 occurs via the respiratory epithelium; high gene expression of *ACE2* and *TMPRSS2* in nasal epithelium^9,10^ have been taken to imply that the nose is a primary entry point for the virus^9^. ACE2 and TMPRSS2 are also co-expressed in bronchial epithelium^9–11^. However, where COVID-19 progresses to severe disease the lung and other organs are also affected. The emerging pattern of severe and fatal COVID-19 disease includes pneumonia with acute respiratory distress syndrome, cytokine storm, widespread vasculopathy, thrombosis, renal failure, hypertension and endothelial dysregulation seen across multiple vascular beds and organ systems^12,13^. Furthermore, COVID-19 is associated with an increased risk of arterial thrombosis^14,15^ and venous thromboembolism with pulmonary embolism^16,17^, both likely to represent an important source of acute and post-COVID-19 morbidity and mortality. While hypertension and thrombosis are common features after COVID-19^18,19^, the important question as to whether COVID-19 as an independent risk factor for cardiovascular disease in the acute setting and during the recovery period, is a concern and remains to be established. Similarly the rates of post-COVID-19 pulmonary fibrosis remain unknown but at discharge the vast majority of patients with COVID-19 have evidence of persisting pulmonary infiltrates on computerised tomography scans^20^ and almost half have physiological impairment^21^. This secondary thrombotic/vascular clinical syndrome of severe COVID-19 suggests that SARS-CoV-2 infects not only respiratory epithelium but also the endothelium disrupting barrier function and allowing access to cardiovascular tissues and other organs of the body^22^. This idea is supported by reports showing that SARS-COV-2 can infect endothelial cells in vitro^23^ and that coronaviruses including SARS-CoV-2 can progress to a systemic infection^24,25^ with some patients showing detectable viral RNA in blood samples ^26–28^.

The reasons that underpin progression of mild to severe or fatal COVID-19 disease remain incompletely understood but risk factors have been defined^29^; these include established cardiovascular disease, diabetes, obesity and black and minority ethnicity [BAME]. However, the dominant risk factor for severe COVID-19 across all datasets is age, with the vast majority of those in hospital with COVID-19 disease being over 40 years. Indeed, a recent disparities in outcomes report by Public Health England found age to be the largest disparity with likelihood of death in adults increasing in an age-dependent manner from around 40 years^30^. Importantly, while positive tests for SARS-CoV-2 infection increase with age the relative rate of infection between adult age groups profoundly underpredicts the effect of advancing age on the risk of death from COVID-19^29,30^. Therefore, understanding how SARS-CoV-2 causes severe disease with pulmonary, thrombotic, cardiorenal and vascular complications is critically important in managing the pandemic and identifying therapeutic strategies.

While some studies report expression profiles of *ACE2* and *TMPRSS2* in epithelial cells^9,11^ and immune cells^10,11^, expression patterns of a wider range of host SARS-CoV-2 entry and processing genes in these cells has was recently reported^11^. However, the relative expression levels of SARS-CoV-2 entry and processing genes in vessels and in endothelial cells has not been fully established. Finally, the impact of age on the expression of these genes in a cardiovascular setting is incompletely understood.

Here we have used publicly available gene expression data to determine the relative expression of key SARS-CoV-2 host entry/ processing genes in human cardiovascular tissues including aorta, coronary artery, heart (atria and left ventricle), whole blood and the kidney and for comparison the colon, spleen and lung. We went on to investigate gene expression in endothelial cells and, for comparison, airway (nasal and bronchial) epithelium and leukocytes (peripheral blood mononuclear cells; PBMCs). We used blood outgrowth endothelial cells as a model because, since they are obtained from blood samples of living donors, data sets across age ranges have been created. Furthermore, blood outgrowth endothelial cells are an accepted model for application in personalised medicine since they retain elements of disease phenotype across a number of cardiovascular and other conditions^31–33^. After mapping gene expression across our target tissues and cells, our primary objective was to determine how age, as the single most dominant risk factor for severe COVID-19, impacts on expression of SARS-CoV-2 entry and processing genes in human cardiovascular and other tissues.

## Methods

### Genotype-Tissue Expression (GTEx) analysis

The Genotype-Tissue Expression (GTEx) project^34^ is an ongoing effort to build a comprehensive public resource to study tissue-specific gene expression and. We downloaded gene expression data from GTEx version 8 (https://www.gtexportal.org/home/datasets) which contain expression data from 54 tissues from 948 donors. We identified tissues of interest based on organ systems affected by severe COVID-19 disease and extracted expression data specifically from those tissues. Tissues were split into two categories; (i) cardiovascular tissues including aorta, coronary artery, heart (atrial and appendage), left ventricle, kidney (cortex) and whole blood and (ii) ‘other tissues’ including lung, colon and spleen. We performed principle component analysis (PCA) on gene expression data from each tissue of interest. We observed that the major variation in gene expression was due to type of death (Hardy Score; Supplementary Figure 1) and so corrected for this. We normalised the gene expression data for each tissue separately using COMBAT-seq^35^ with Hardy score as a batch. After normalisation expression data was extracted for our target genes (*ACE, ACE2, ADAM17, BSG, CTSB, CTSL, FURIN, PPIA, PPIB* and *TMPRSS2*). The following number of donors were identified for each tissues; aorta (432), coronary artery (240), atrial appendage (429), left ventricle (432), kidney cortex (85), whole blood (755), lung (578), colon (779) and spleen (241). Age identifiers in GTEx are grouped by decade, as such results were analysed based on samples that associated with 20-29 to 70-79 years of age. Principle component analysis (PCA) plots of raw and processed GTEx data is presented in Supplementary Figure 1.

### Gene expression dataset systematic review analysis

Using ArrayExpress and NCBI GEO we identified the raw datasets (.CEL files) of transcriptomic gene expression profiling by microarray of healthy adult donors for blood outgrowth endothelial cells, peripheral blood mononuclear cells (PBMCs) and bronchial airway (obtained from bronchial brushing) and nasal (obtained nasal brushing) epithelium. We applied strict inclusion criteria; (i) only datasets that used Affymetrix Gene Chips (.CEL files) were included, only datasets where individual ages are defined were included and (iii) only datasets for ‘untreated’ cells were included. The following studies and number of donors were identified (see Supplementary Table 1); blood outgrowth endothelial cells; 3 studies with 63 donors, PBMCs; 6 studies with 84 donors, airway (bronchial) epithelium; 2 studies with 74 donors and nasal epithelium; 3 studies with 111 donors.

### Transcriptomic expression profiling of cell data

Raw (.CEL) files were imported into Partek Flow^®^ software and aligned with STAR to the human assembly (hg19) whole genome. The data was quantified to an annotation model using ‘Ensembl Transcripts release 75’ and normalised to ‘Counts Per Million’ and filtered to remove genes below the reliable quantitation threshold. The gene expression values from different studies were merged based on gene names. To correct for the batch effects, the data was normalised using the empirical Bayes model ComBat^36^. The normalised expression values of our target genes (*ACE2, CYPA, CYPB, BSG, ADAM17, TMPRSS2, FURIN, CTSB* and *CTSL*) were then extracted for further downstream analysis. PCA plots of raw and processed cell data is presented in Supplementary Figure 1.

### Statistical analysis

All data were analysed on GraphPad Prism v8 and are shown as mean +/− SEM for samples from ‘n’ = individual donors. Data were grouped into two groups; samples from adults below the age of 40 and above the age of 40. Data were tested for normality of distribution and analysed using parametric (Student T-test) or non-parametric (Mann Whitney U-test) tests as appropriate as a discovery exercise. Where p<0.05 follow on correlation tests were performed using Pearson’s, for continuous variables (cell data), or Spearman’s, for ordinal variables (tissue data), tests. Details of tests used are given in individual figure legends.

## Results

We quantified nine SARS-CoV-2 entry and processing genes (*ACE2*, *BSG*, *ADAM17*, *TMTRSS2, CYPA, CYPB, CTSB, CTSL* and *FURIN*) (Figure 1 and Figure 2) along with *ACE* in cardiovascular tissues (including kidney and whole blood) and other organs (including lung spleen and colon), (Figure 1) and in endothelial cells, respiratory epithelial cells and in PBMCs (Figure 2). ACE is not directly related to cellular processing of the virus but represents a pharmacological link with ACE2.

**Figure 1:**
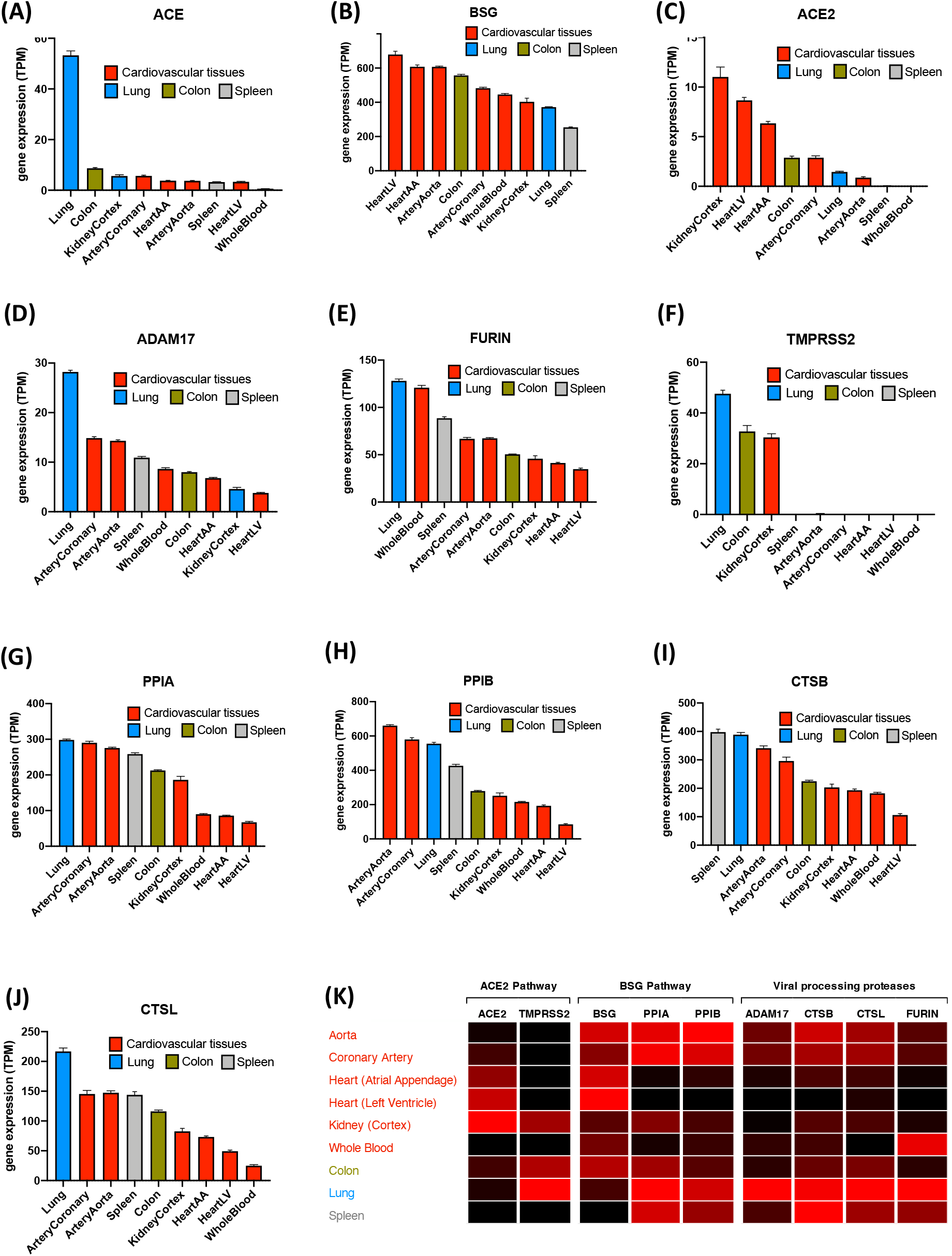
Expression of SARS-CoV-2 entrance/processing genes in different cardiovascular and non-cardiovascular tissues. Standardised expression levels for the genes *ACE(A), BSG (B), ACE2 (C), ADAM17 (D), FURIN (E), TMPRSS2 (F), PPIA (G), PPIB (H), CTSB (I) and CTSL (J)* were obtained from GTEx from cardiovascular tissues (aorta, coronary artery, atrial appendage, left ventricle, kidney cortex, and whole blood; red columns) and other tissues (lung, blue columns; spleen, grey columns and colon, green columns). Data for each tissue was corrected for batch effects COMBAT-seq and expressed as mean +/− S.E.M. Tissues were ranked in order of expression for each gene. A heat map showing expression of ACE2 and BSG pathways and viral processing proteases in each tissue was generated (K). Data were coloured by gene, whereby black is the lowest expressing tissue and red is highest expressing tissue.

**Figure 2:**
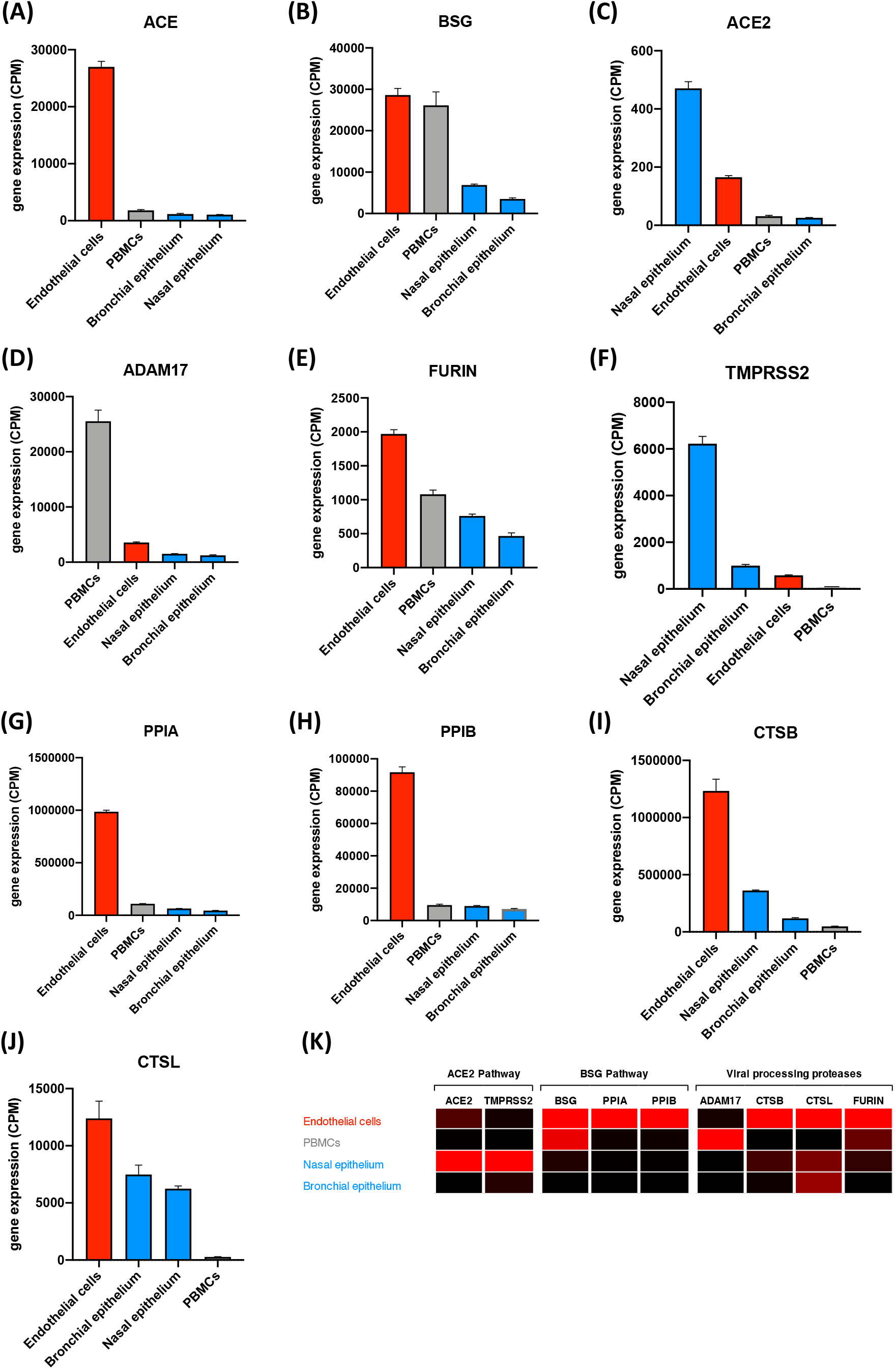
Expression of SARS-CoV-2 entrance/processing genes in blood outgrowth endothelial cells, PBMCs and epithelial cells (nasal and bronchial) Standardised expression levels for the genes *ACE(A), BSG (B), ACE2 (C), ADAM17 (D), FURIN (E), TMPRSS2 (F), PPIA (G), PPIB (H), CTSB (I) and CTSL (J)* were obtained from online databases from human blood outgrowth endothelial cells (Endothelial cells, red columns), peripheral blood mononuclear cells (PBMCs, grey columns) and epithelial cells (nasal and bronchial, blue columns). The data was aligned and analysed using PartekFlow^®^ and corrected for batch effects using COMBAT-seq and expressed as mean +/− S.E.M. Cells were ranked in order of expression each gene. A heat map showing expression of ACE2 and BSG pathways and viral processing proteases in each cell type was generated (K). Data were coloured by gene, whereby black is the lowest expressing cell type and red is highest expressing cell type.

### Relative expression of SARS-CoV-2 entry genes across organs (Figure 1 and Figure 3)

**Figure 3:**
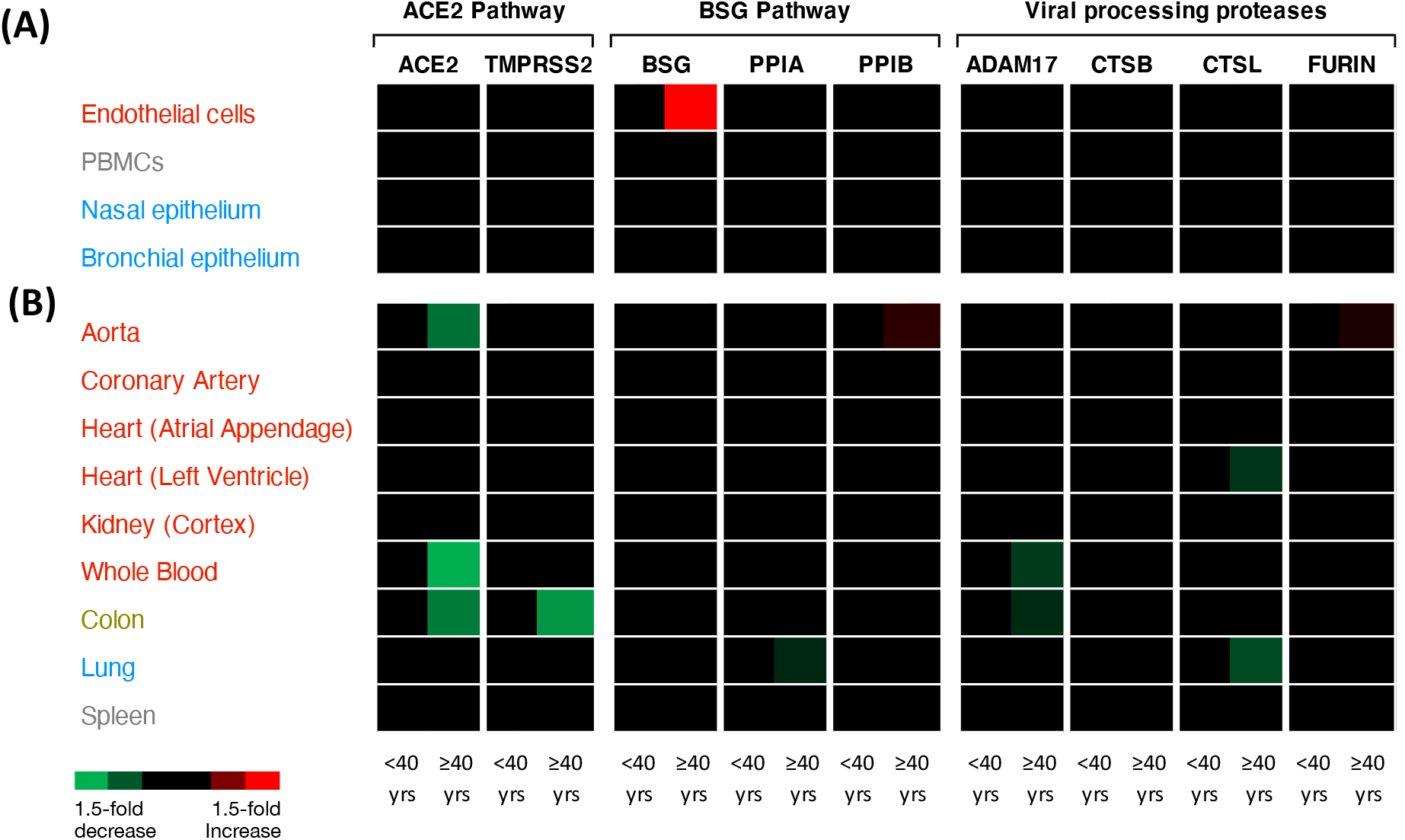
Heat map representing expression of SARS-CoV-2 entrance/processing genes significantly altered by age. Heatmaps were generated for the expression of *ACE, ACE2, BSG, ADAM17, CTSB, CTSL, FURIN, PPIA, PPIB and TMPRSS2* in cells (A) and organs (B). Data were analysed based on two adult age groups; under or over 40 years. Data were analysed using an unpaired Mann-Whitney t-test or unpaired t-test depending on normality distributions. Significant data (p<0.05) are shown as either increased (red) or decreased (green) expression; black corresponds to no significant change.

As expected, *ACE* was highly expressed in the lung^37^ with lower but consistent levels expressed across other tissues and with very low levels present in blood. Of the two putative SARS-CoV-2 receptors, *BSG* was highly expressed across all tissues with higher levels seen in most cardiovascular tissues than in the lung or spleen. *ACE2*, across all tissues, was expressed in relatively low levels. However, cardiovascular tissues including kidney, heart and blood vessel (coronary artery) expressed higher levels of *ACE2* than the lung or spleen. Relatively low levels of *ACE2* were seen in whole blood. The colon was positioned mid group for both *BSG* and *ACE2* expression. Of the putative processing genes, required for spike protein conditioning and/or cleavage of *ACE2* allowing viral entry, *ADAM17* and *FURIN* were each enriched in the lung with relatively stable levels of expression across cardiovascular and other target tissues. *TMPRSS2* was also enriched in the lung, colon and kidney cortex with very low levels present in arteries, heart, spleen and blood. For the putative vial partner ligands of *BSG*, *PPIA* and *PPIB,* both were expressed throughout our selected tissues with higher levels expressed in arteries than kidney, blood or heart tissues. Lung, spleen and colon expressed high or midranking levels of *PPIA* and *PPIB*. The endosomal proteases, *CTSB* and *CTSL* showed similar expression patterns across our tissues of interest. Both *CTSB* and *CTSL* were enriched in the lung and spleen (*CTSB*) with relatively high levels across all tissues. *CTSB* and *CTSL* were more highly expressed in arteries than kidney, heart or blood.

### Relative expression of SARS-CoV-2 entry genes in endothelial cells versus airway epithelium and PBMCs (Figure 2 and Figure 3)

To complement and extend the above organ-level approach we focussed on endothelial cells versus respiratory epithelial cell types and immune cells (PBMCs), in line with COVID-19 disease pathology. As expected, *ACE* was highly enriched in endothelial cells^38^ with lower levels present in PBMCs, bronchial and nasal epithelium. *BSG* was enriched in endothelial cells and PBMCs with lower levels expressed in nasal and bronchial airway epithelium. *ACE2* was enriched in nasal epithelium flowed by endothelial cells and lower levels in PBMCs and bronchial epithelium. *ADAM17* was highly enriched in PBMCs followed by endothelial cells and nasal and bronchial epithelial cells. *FURIN* was enriched in endothelial cells with mid-ranking levels expressed in PBMCs and lower levels in nasal and bronchial epithelium; *TMPRSS2* was highly enriched in nasal epithelial cells followed by bronchial epithelial cells, endothelial cells and PBMCs. *PPIA* and *PPIB* were highly expressed in endothelial cells with lower levels in PBMCs, nasal and bronchial epithelial cells. Intracellular proteases *CTSB* and *CTSL* were both also enriched in endothelial cells followed by airway epithelium (*CTSB*, nasal>bronchial; *CTSL*, bronchial>nasal) and low levels in PBMCs.

Next, in line with recent Public Health England’s review of disparities in risks and outcomes for COVID-19^17^, we grouped data into two age categories, <40 and >40 years to determine differences in gene expression. Where differences were found and based on clinical evidence showing that the risk of death from COVID-19 directly correlates with age^19^, we performed follow-on correlation analysis.

### Effect of age on expression of SARS-CoV-2 entry genes across cardiovascular and COVID-19 target tissues

#### Arteries

In the aorta, *FURIN and PPIB* were increased while *ACE2* was deceased in samples from adults >40 years of age (Figure 3). Reductions of *ACE2* or increases of *PPIB* linearly correlated with age (Figure 4; Supplementary Figure 2). *FURIN* expression did not linearly correlate with age (Supplementary Figure 2). None of our selected genes were affected by age in the coronary artery (Figure 3).

**Figure 4:**
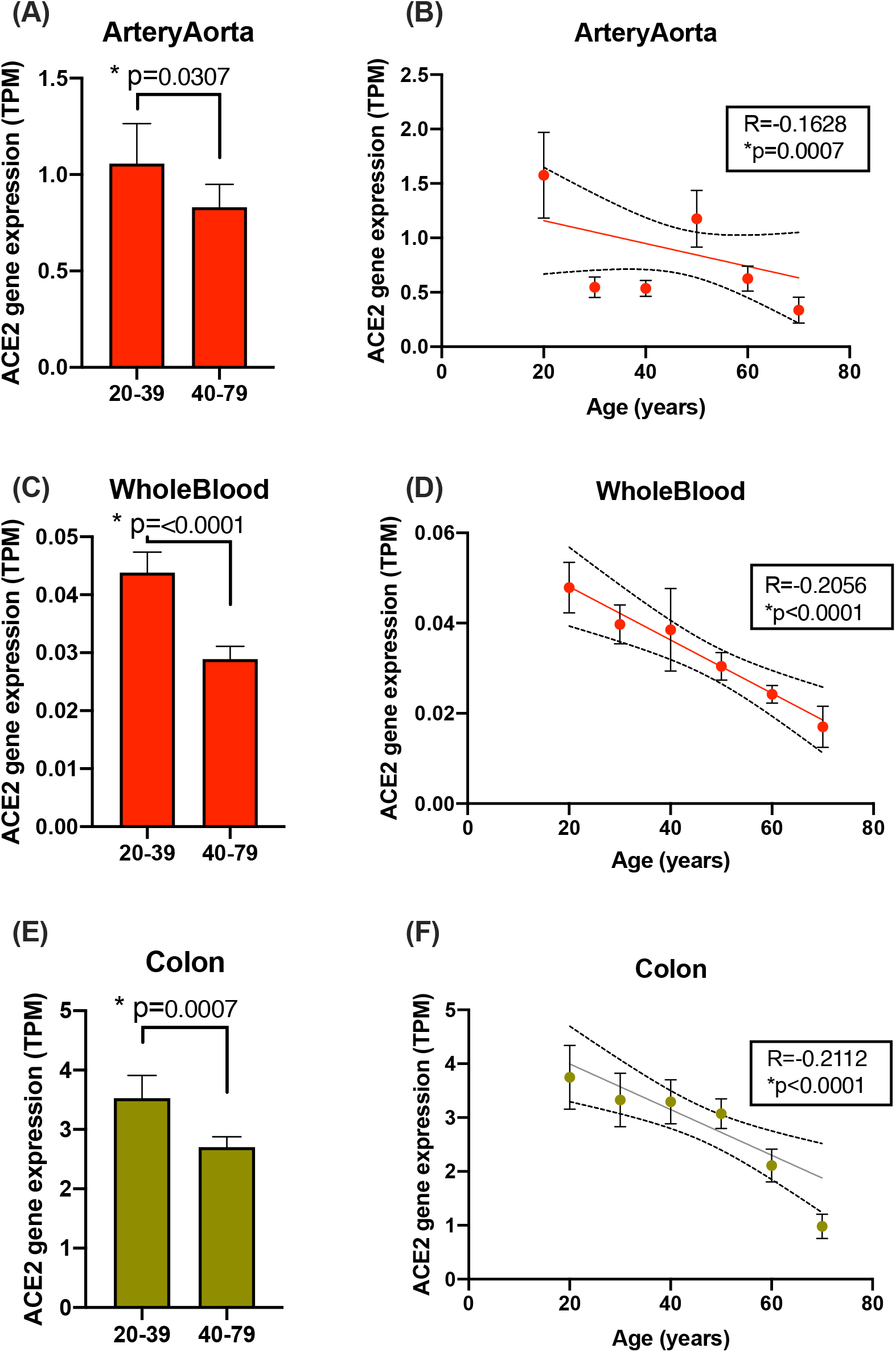
The effect of age on *ACE2* expression in aorta (A,B), whole blood (C,D) and colon (E,F) *ACE2* levels in aorta (A,B), whole blood (C,D) and colon (E,F) were analysed based on adults under vs over 40-year-olds (A, C, E). Data is expressed as mean +/− S.E.M. and analysed using unpaired Mann-Whitney T-test (A,C,E) or Spearman’s correlation test (B,D,F); significance was accepted when *p<0.05.

#### Heart and Kidney

In the heart (atria and left ventricle) or kidney none of our selected genes were affected by age (Figure 3).

#### Whole Blood

In whole blood *ACE* expression increased while *ACE2 and ADAM17* were reduced in samples from individuals >40 years of age (Figure 3; Figure 4; Supplementary Figure 4). Of these genes reduced expression of *ACE2* (Figure 4) and *ADAM17* and increased expression of *ACE* linearly correlated with age (Supplementary Figure 4).

#### Colon

In the colon *ACE2, ADAM17* and *TMPRSS2* were decreased in samples from individuals >40 years of age (Figure 3; Figure 4; Supplementary Figure 5). Of these genes, reduced expression levels of *ACE2* and *TMPRSS2* but not *ADAM17* linearly correlated with age (Supplementary Figure 5).

#### Lung

In the lung expression levels of *CTSL* and *PPIA* were decreased in samples from adults >40 years of age (Figure 3). Reduced expression linearly correlated with age for *PPIA* but not *CTSL* (Supplementary Figure 6).

#### Spleen

None of our studied genes were altered with age in the spleen (Figure 3).

### Effect of age on expression of SARS-CoV-2 entry genes in endothelial cells, airway cells and leukocytes

In endothelial cells *BSG*, but not other genes, was increased in samples from adults >40 years (Figure 3 and Figure 5; Supplementary Figure 7) and levels showed a positive linear correlation with age (Figure 5). By contrast, only *ACE* was fractionally (but statistically significantly) reduced in nasal epithelium between age categories (Figure 3; Supplementary Figure 8) and this did not linearly correlate with age (Supplementary Figure 9). No genes were altered in bronchial epithelium (Supplementary Figure 10) and PBMCs (Supplementary Figure 11) with age (Figure 3).

**Figure 5:**
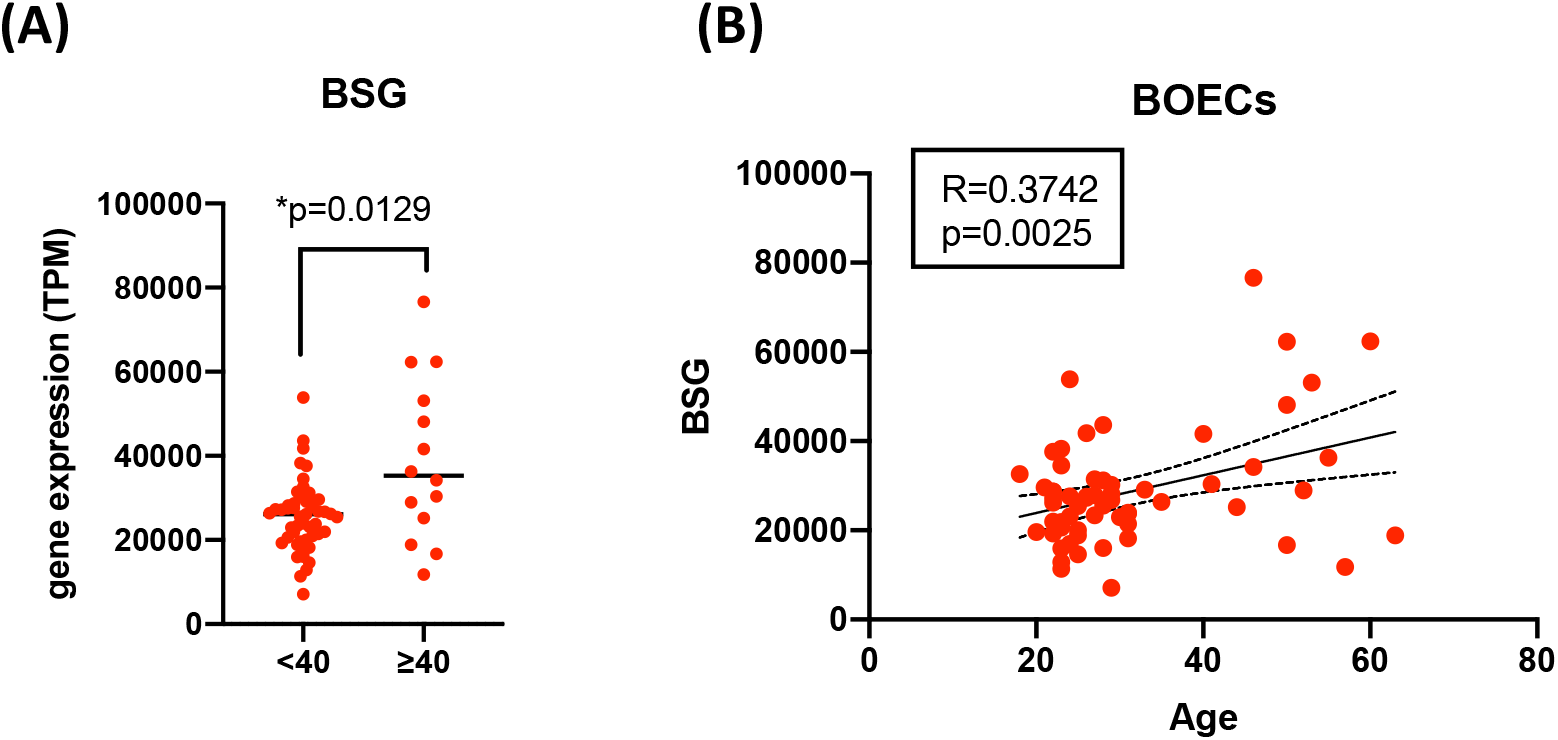
The effect of age on *BSG* expression in blood outgrowth endothelial cells. *BSG* levels in blood outgrowth endothelial cells (Endothelial cells) were analysed in adults under 40 years (<40) versus over 40 years (>40) using an unpaired Mann-Whitney T-test (A) and correlations with age determined using Pearson’s correlation analysis (B); significance was accepted when *p<0.05.

### Summary

In light of the cardiovascular/thrombotic sequalae associated with severe COVID-19 disease and the overwhelming risk that increased age carries, our aim was to obtain mechanistic insight by interrogating gene expression profiles in cardiovascular tissues and cells. Our focus was on the two putative receptors for SARS-CoV-2, *ACE2* and *BSG* along with a selected range of genes thought to be involved in virus binding/processing. In this study we have made four important observations: (i) Cardiovascular tissues and/or endothelial cells express the required genes for SARS-CoV-2 infection, (ii) SASR-CoV-2 receptor pathways, *ACE2*/*TMPRSS2* and *BSG*/*PPIB(A*) polarise to lung/epithelium and vessel/endothelium respectively, (iii) expression of SARS-CoV-2 host genes are, on the whole, relatively stable with age and (iv) notable exceptions were *ACE2* which decreases with age in some tissues and *BSG* which increases with age in endothelial cells.

### Discussion

Initial SARS-CoV-2 infection occurs in the airways. For most people infection is either asymptomatic or associated with mild symptoms consistent with localised viral infection in respiratory tissues. Naturally, therefore, most research to date has focused on investigations in the lung or airway cells and understanding how to manage the complications of pneumonia and ventilation failure. However, in some individuals SARS-CoV-2 infection progresses to severe COVID-19 disease, which is a systemic illness with complications specifically associated with the cardiorenal system, endothelium and thrombosis. Now, understanding the vascular component of severe COVID-19 disease associated with SARS-CoV-2 infection is emerging as an urgent unmet clinical need.

Here, we report that SARS-CoV-2 receptors and processing genes are expressed across all cardiovascular target tissues and/or in endothelial cells, supporting the idea that systemic organs contain the required machinery to be infected by the virus. In regard to the two SARS-CoV-2 receptor pathways; expression levels of *ACE2* were higher in cardiovascular tissues than the lung. By contrast, *TMPRSS2*, thought to be required for SARS-CoV-2 infection in epithelial cells, was present in lung colon and kidney but essentially absent in other cardiovascular tissues (heart, vessels and whole blood). Our findings describing the relative levels of these genes in human tissues are in line with others using similar approaches for *ACE2*^39–43^ and *TMPRSS2*^41,44^. Moreover, we found that both *ACE2* and *TMPRSS2* were enriched in nasal epithelium with low levels in bronchial epithelium and PBMCs which is in agreement with recent work from others (nasal versus bronchial epithelium^9,10^ and versus PBMCs^10^). Radzikowsa et al., profiled a wider range of SARS-CoV-2 entry genes in immune cells and differentiated primary bronchial epithelial cells and also reported relatively high levels of expression of *PPIA, BSG, PPIB* with much lower levels of *TMPRSS2* followed by *ACE2* in airway cells^11^. However, our focus was on the cardiovascular system. Importantly we found endothelial cells to express much lower levels of *ACE2* and *TMPRSS2* than nasal epithelium but higher levels than those expressed in PBMCs. Nevertheless, in contrast to the *ACE2/TMPRSS2* pathway genes, *BSG* and *PPIB* were expressed in higher levels in vessels and endothelial cells than in the lung and airway epithelial cells. Together our findings suggest that SARS-Cov-2 and other relevant viruses exploit BSG as a receptor pathway in the vasculature. Our findings align with recent work by Ganier and colleagues^45^ and add evidence to the recent ‘proposed mechanism’ explaining how SARS-CoV-2 accesses endothelial cells, presented by Acosta Saltos and Acosta Saltos^46^.

Severe COVID-19 is exceptionally rare in children. In adults the strongest risk factor for severe disease and death is age, with those under 40 years being at very low risk; the risk of severe COVID-19 disease and death increases proportionally after the age of 40^30^. Of the genes that we studied, several candidates, including *ACE2*, were affected by age but with the exception of *BSG* in endothelial cells and *PPIB* and *FURIN* in aorta, expression was reduced in those >40. We found consistent age-related reductions in *ACE2* in whole blood, aorta and in the colon. Our findings are in line with those published by Chen and co-workers who also reported a negative correlation between *ACE2* and age in a range of tissues including colon and blood^39^. Moreover, our work corroborates earlier studies showing that ACE2 (protein) declines with age in mouse aorta^47^. Other studies in rats also showed that ACE2 declines with age in the lung and kidney^48,49^. It should be noted, however, that Li and colleagues found no effect of age on *ACE2* expression across a similar selection of tissues^43^ and that Santesmasses and colleagues found that *ACE2* expression increased with age in the lung^50^. We also found a trend for *ACE2* to increase in the lung but this did not reach statistical significance in our study. Key differences between the studies include the analytical approaches applied, the number of tissues selected, and the age groups used.

Since ACE2 is a receptor for SARS-CoV-2, which declines with age in some settings (this study)^43,47–49^ and because age is the strongest predictor for fatal COVID-19 disease a paradox has emerged^51^: *SARS-CoV-2 receptor expression does not positively correlate with high risk groups of severe COVID-19 disease*. One explanation of the paradox has been that because ACE2 is a cardioprotective enzyme, while ACE2 is the receptor for airway infection of SARS-CoV-2, low levels of ACE2 in the circulation of the elderly and those with cardiovascular disease increases the risk of cardiovascular complications associated with severe COVID-19 disease^51^. Additionally, it has been hypothesised that BSG acts as a receptor for SARS-CoV-2 in endothelial cells^46^. BSG expression is increased in a range of cardiovascular diseases which could compensate for any age/disease associated reductions in ACE2 in regard to viral infection. In line with this in our study, *BSG* positively correlated with age and this association was only seen in endothelial cells. Others have found that BSG increases with age in the skin^52^. It is not clear why we did not see age related increases in *BSG* in the aorta or coronary artery or in organ samples. One explanation could be that in vessels the delicate lining of the endothelium may have been lost during tissue dissection and/or that the age effects on *BSG* expression in endothelial cells is diluted out in complex tissues by expression levels in other cells which make up the bulk of the samples. It should also be considered that whilst blood outgrowth endothelial cells display key ubiquitous features of endothelial cells and retain disease phenotypes^31^, heterogeneity exists within endothelial cells as they age with passage and in different vascular beds.

Based on our findings together the cardiovascular, renal and thrombotic complications associated with severe COVID-19 disease, we suggest that BSG expression in the pulmonary and systemic vasculature maybe an important driver which explains the heightened risk of severe disease and death observed in those >40 years of age. These observations add to the growing evidence and provide additional mechanistic insight supporting the targeting the BSG: PPIA/PPIB axis in severe COVID-19 disease.

BSG is an attractive lead to have emerged from this work. BSG is upregulated in a range of diseases including those co-morbidities/morbidities associated with increased risk of severe COVID-19 disease and poorer outcomes including thrombosis^53^, pulmonary hypertension^54^, renal disease^55^, obesity^11^ and diabetes^56^. Moreover BSG, as both a receptor for SARS-Cov-2 and as an inducer of extracellular matrix metalloproteinases, may be relevant when considering the potential role of the vasculature and, specifically endothelial dysfunction, in propagating and driving pulmonary fibrosis^57,58^ – one of the most feared long term complications of COVID-19^59^.

Clearly, a full understanding of BSG interactions with SARS-CoV-2 will provide valuable mechanistic insight and could identify new therapeutic targets and/or provide additional insight for experimental drugs currently in trials for the prevention and treatment of severe COVID-19 disease. For example, cyclosporin, an immunosuppressive drug used in the prevention of organ rejection after transplant, has been suggested as a therapy for cytokine storm associated with severe COVID-19 disease. To this end clinical trials to assess cyclosporin drugs in COVID-19 have been initiated^42,43^. Cyclosporin also binds PPIA and therefore may additionally inhibit viral binding to BSG. Our findings that BSG expression is increased with age may provide additional rational for the use of drugs that interfere with cyclophilin binding. In light of our findings and the overwhelming clinical information indicating vascular inflammation in severe COVID-19 disease, the relative role of SARS-CoV-2 receptors and processing proteins in endothelial cells should be investigated. Our data indicates that blood outgrowth endothelial cells will be a useful tool in future work exploring mechanisms of viral infection and inflammation in COVID-19. In addition, since blood outgrowth endothelial cells can be obtained from blood samples of living donors, functional assays using these cells from protected and at-risk populations may provide a means of identifying personalised therapies and those at risk of severe disease.

Since BSG is utilised by other pathogens our findings have implication beyond the current pandemic. Finally, because BSG is implicated in a range of cardiovascular diseases and fibrosis our observations may have relevance to our understanding of the diseases associated with aging.

### Limitations and future studies

The details of the cellular events involved in SARS-CoV-2 infection in different cell types has not yet been fully established which means that the weighted relevance of genes that we have investigated in COVID-19 remains to be elucidated. However, each of the genes that we studied are established in various aspects of human physiology and pathology and so our analysis has relevance to the understanding of aging in a setting wider than infection. We used publicly available data sets which, while were analysed in a systematic and unbiased manner, require replication and validation in a prospective follow up clinical and mechanistic studies. Furthermore, gene expression data invariably requires biological validation in functional assays.

---

**Supplementary Table 1:**
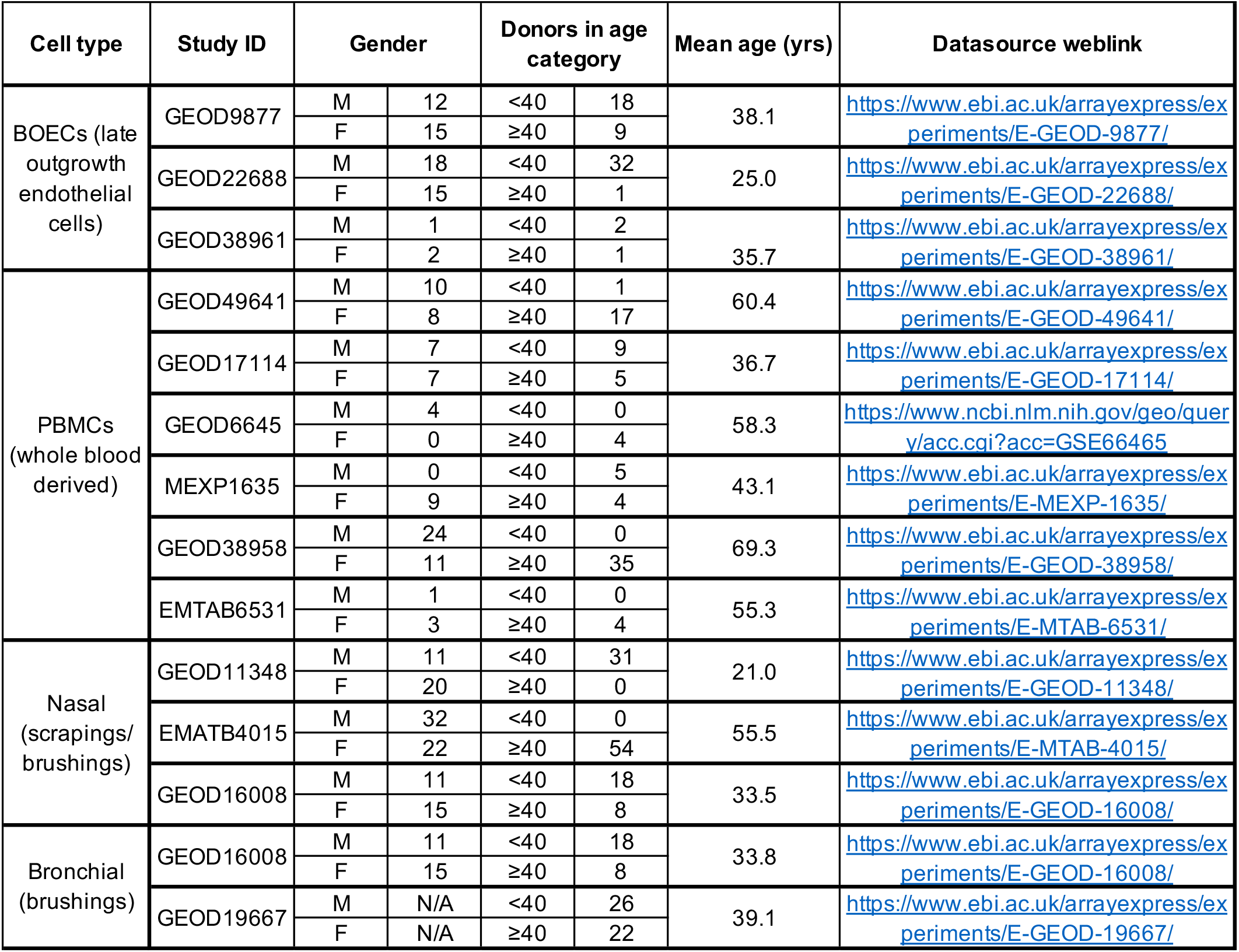
Identified cell studies. Raw transcriptomic gene expression profiling datasets were obtained from ArrayExpress and NCBI GEO for blood outgrowth endothelial cells (BOECs; also known as ‘late outgrowth endothelial cells’), peripheral blood mononuclear cells (PBMCs) and bronchial airway (obtained from bronchial brushing) and nasal (obtained nasal brushing) epithelium.

**Supplementary Figure 1:**
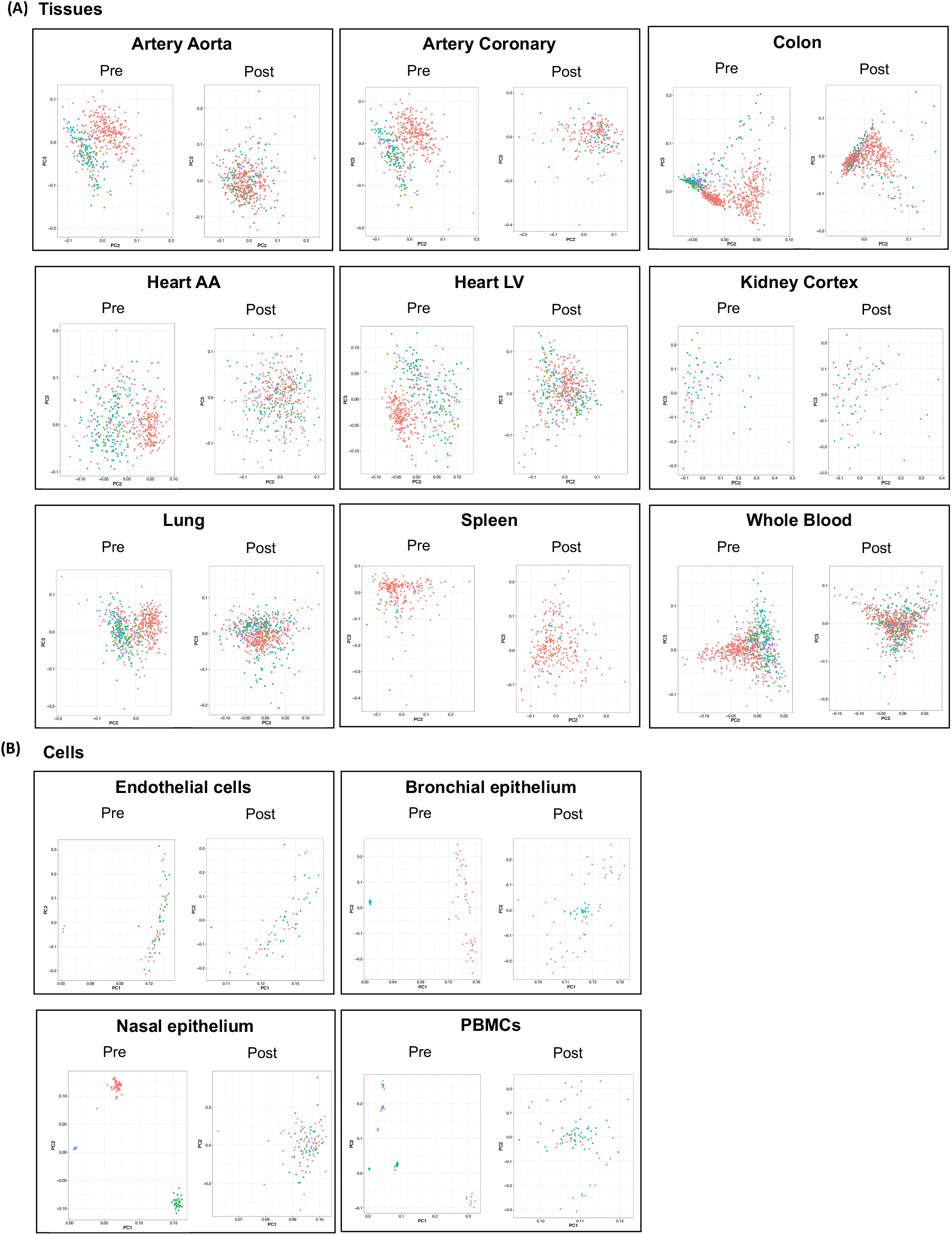
Principle component analysis (PCA) of genes of interest pre and post COMBAT-Seq normalisation. PCA plots for all organs (A) and cells (B) pre and post COMBAT-Seq normalisation were generated.

**Supplementary Figure 2:**
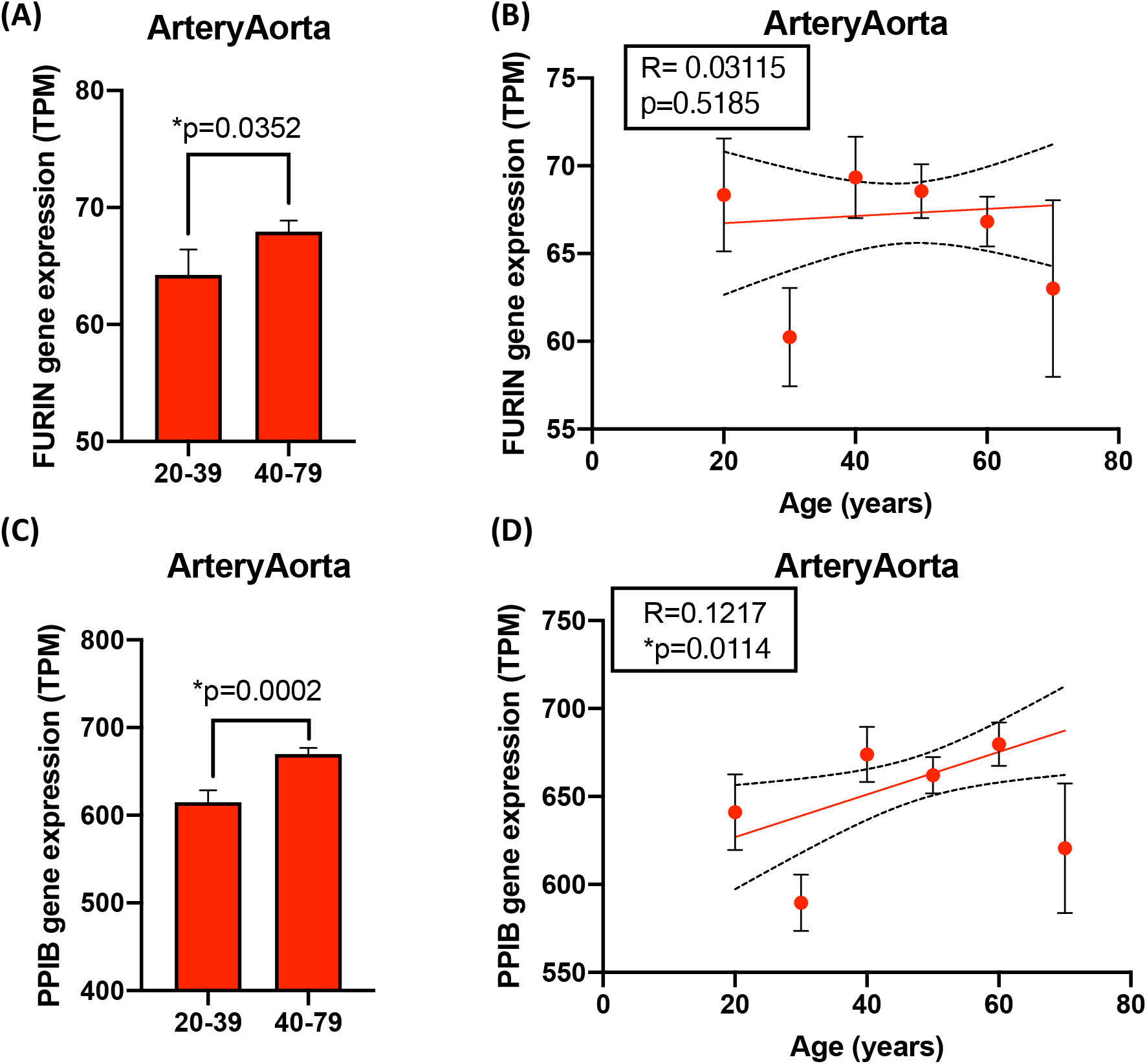
The effect of age on *FURIN* (A,B) and *PPIPB* (C,D) in aorta. Gene expression levels in aorta are shown as mean +/− S.E.M and were analysed based on adults under vs over 40-year-olds using an unpaired Mann-Whitney T-test (A,C) and correlations with age determined using Spearman’s correlation test analysis (B,D); significance was accepted when *p<0.05.

**Supplementary Figure 3:**
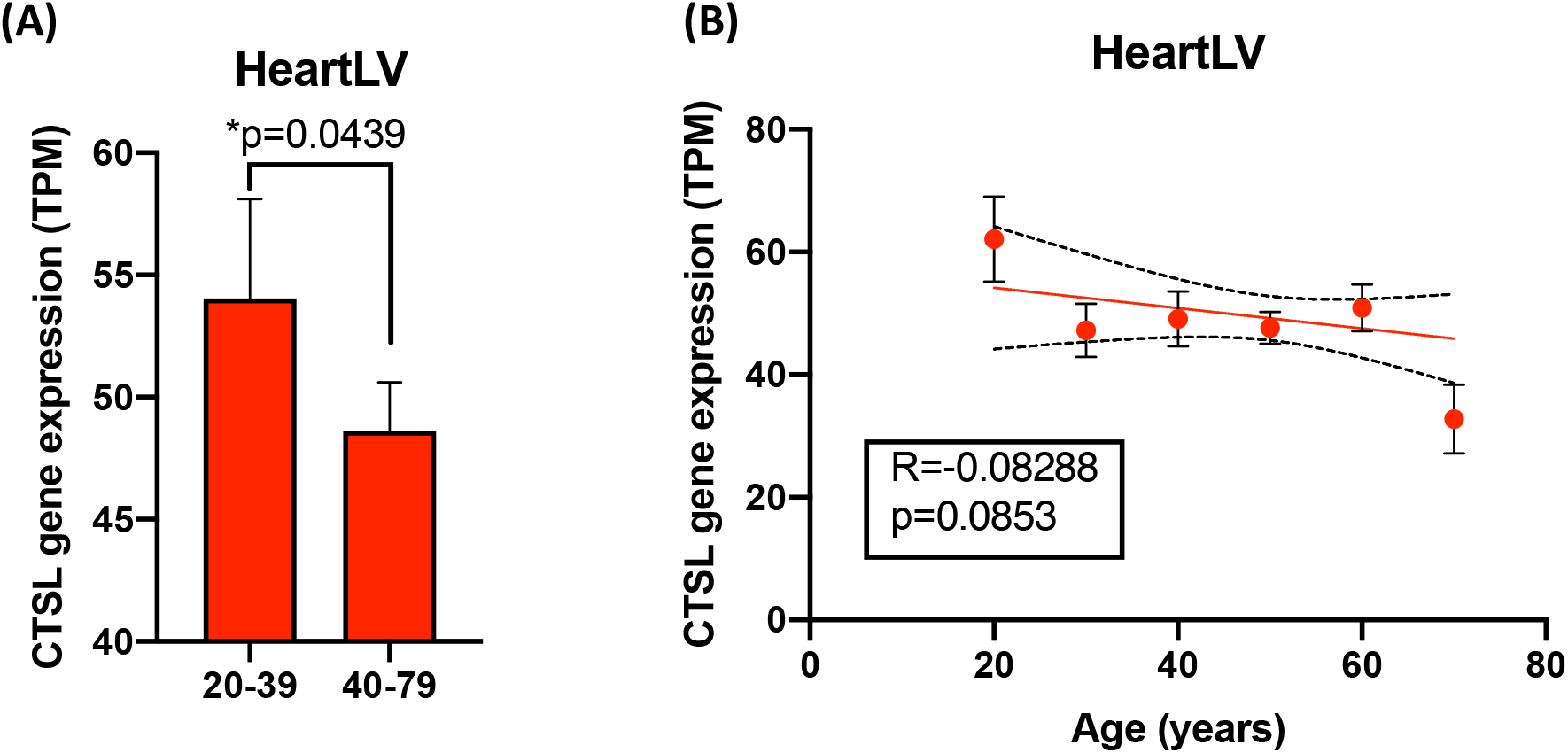
The effect of age *CTSL* heart left ventricle (heartLV) Gene expression levels in heartLV are shown as mean +/− S.E.M and were analysed in adults under 40 years (<40) versus over 40 years (>40) using an unpaired Mann-Whitney T-test (A) and correlations with age determined using Spearman’s correlation test analysis (B); significance was accepted when *p<0.05.

**Supplementary Figure 4:**
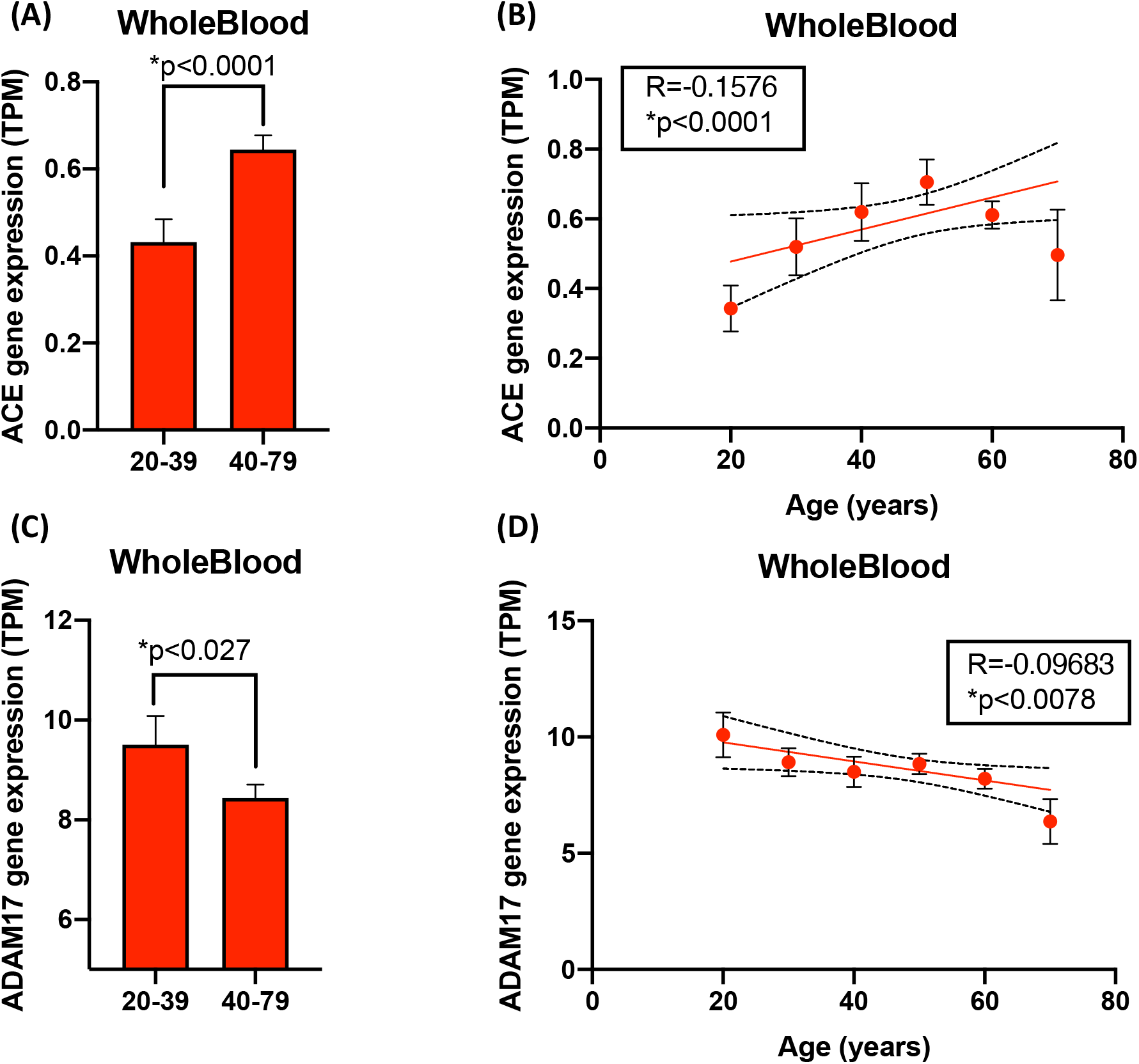
The effect of age on *ACE* (A,B) and *ADAM17* (C,D) in whole blood. Gene expression levels in whole blood are shown as mean +/− S.E.M and were analysed in adults under 40 years (<40) versus over 40 years (>40) using an unpaired Mann-Whitney T-test (A,C) and correlations with age determined using Spearman’s correlation test analysis (B,D); significance was accepted when *p<0.05.

**Supplementary Figure 5:**
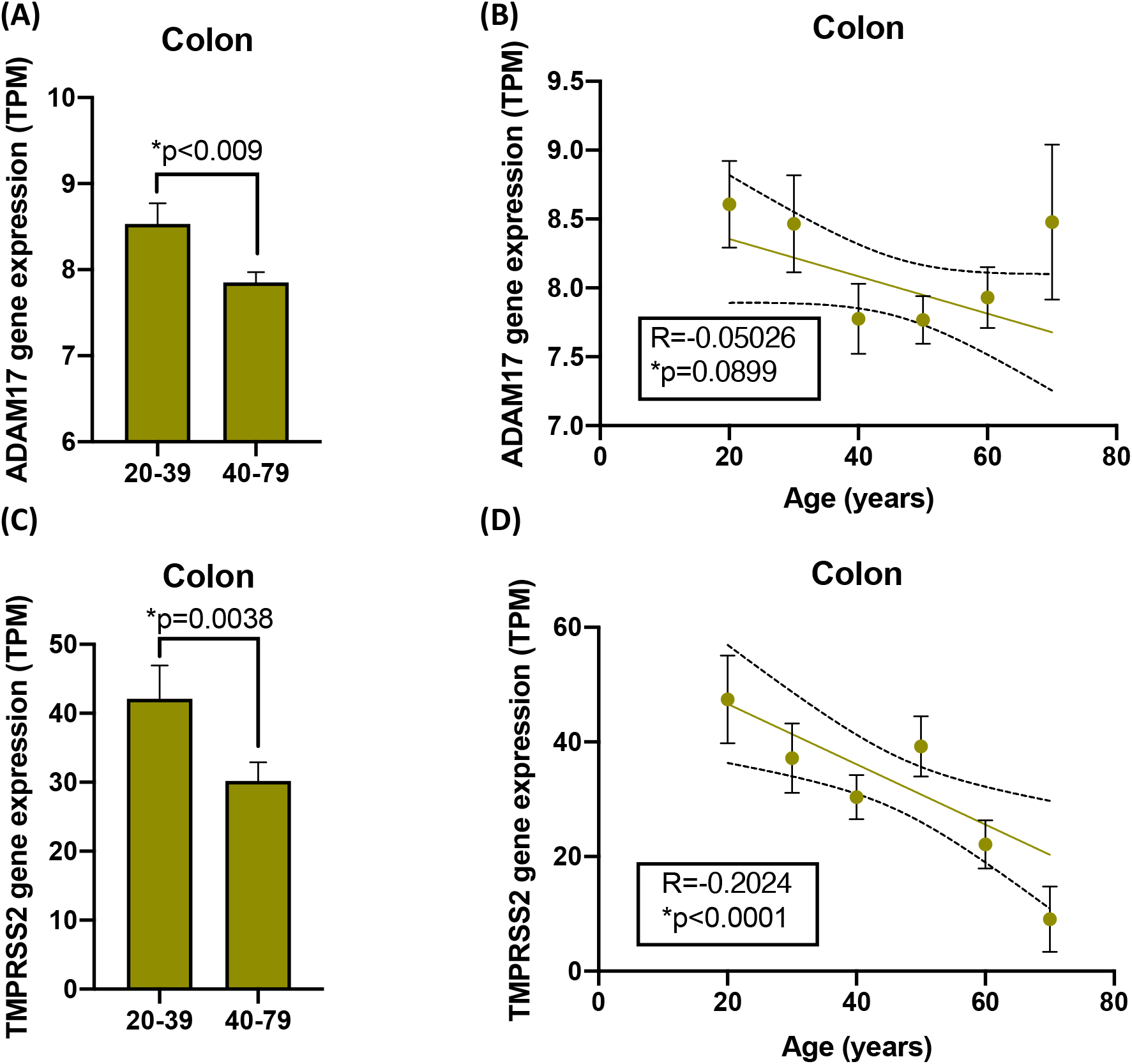
The effect of age on *ADAM17* (A,B) and *TMPRSS2* (C,D) in colon. Gene expression levels in colon were analysed in adults under 40 years (<40) versus over 40 years (>40) using an unpaired Mann-Whitney T-test (A,C) and correlations with age determined using Spearman’s correlation test analysis (B,D); significance was accepted when *p<0.05.

**Supplementary Figure 6:**
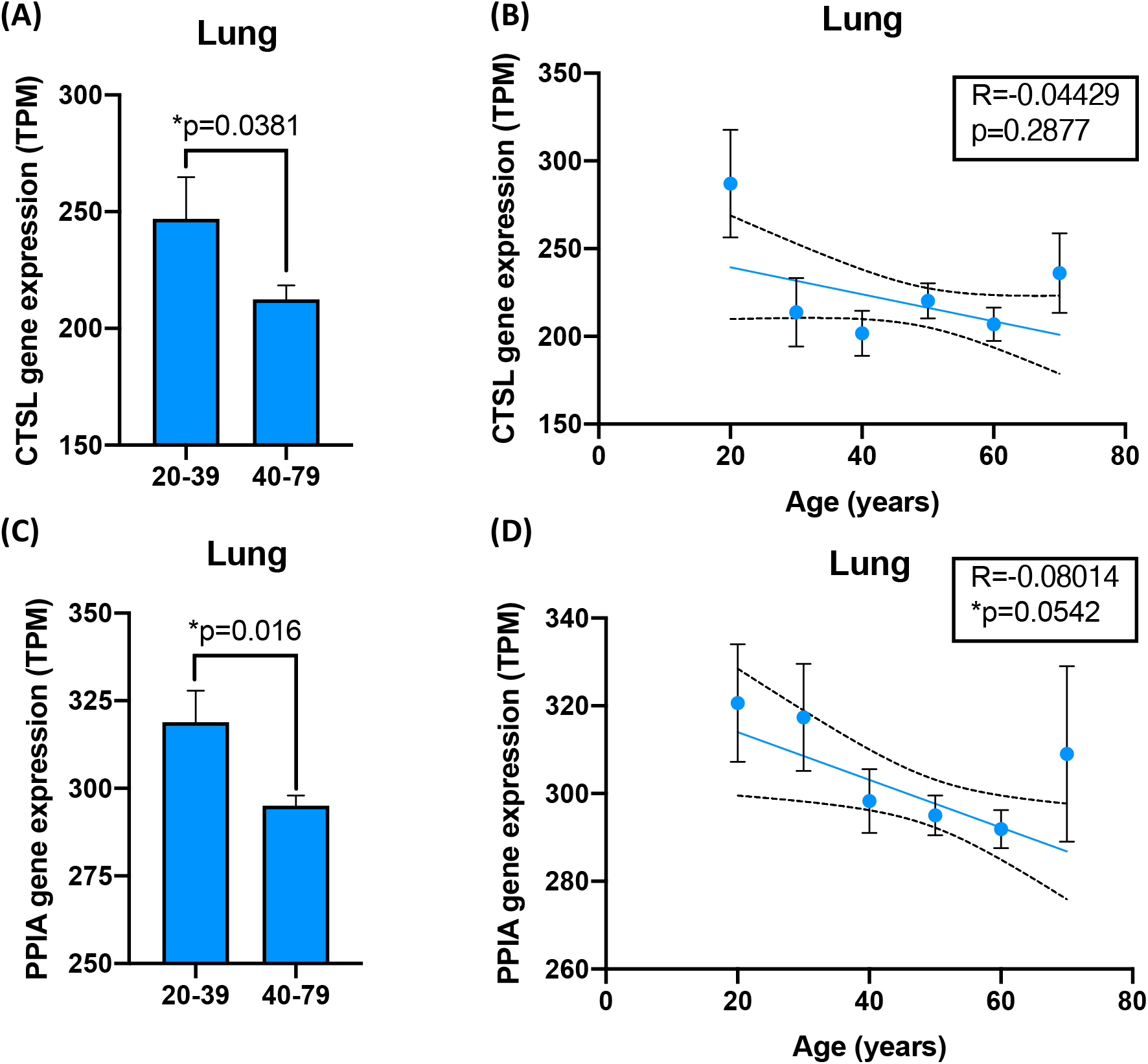
The effect of age on *CTSL* (A,B) and *PPIA* (C,D) in lung. Gene expression levels in lung were analysed in adults under 40 years (<40) versus over 40 years (>40) using an unpaired Mann-Whitney T-test (A,C) and correlations with age determined using Spearman’s correlation test analysis (B,D); significance was accepted when *p<0.05.

**Supplementary Figure 7:**
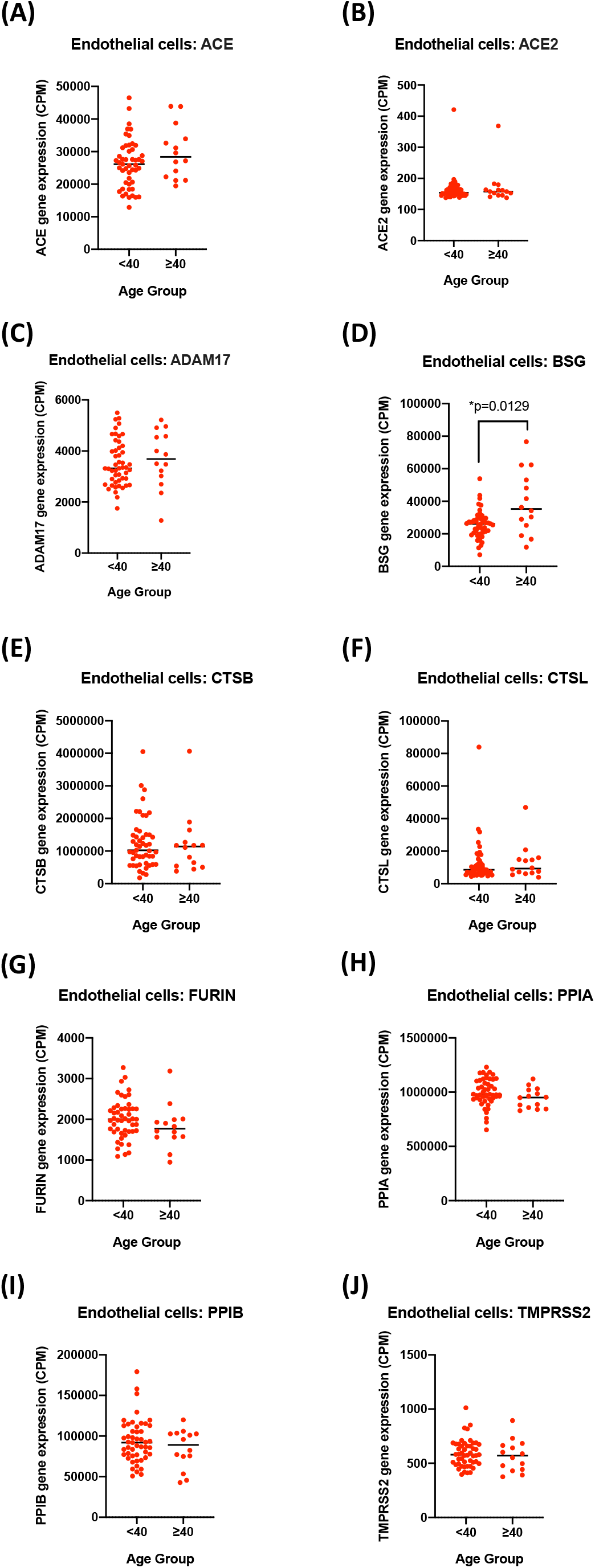
The effect of age (adult under 40 years; <40 versus over 40 years; >40) on of SARS-CoV-2 entrance/processing genes in blood outgrowth endothelial cells. Data is presented as individual points for each gene and analysed using Students T-Test or Mann-Whitney T-test as appropriated; significance was accepted when *p<0.05.

**Supplementary Figure 8:**
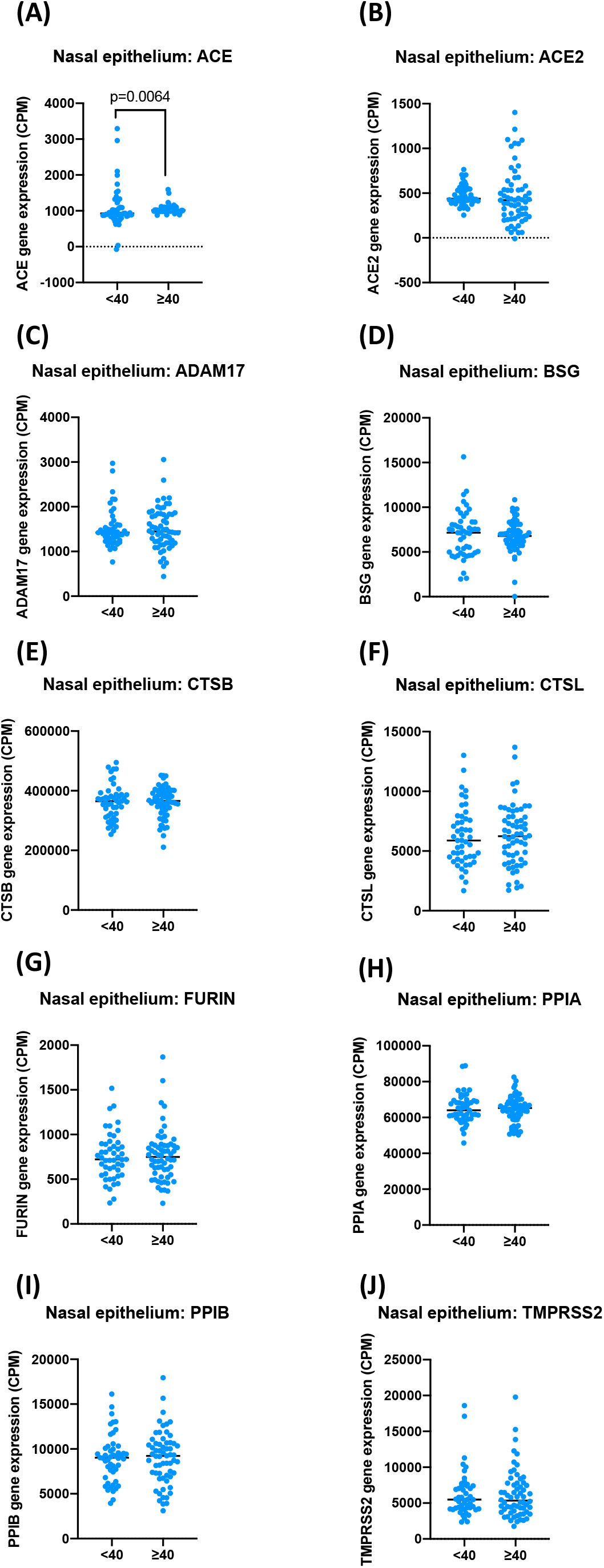
The effect of age (adult under 40 years; <40 versus over 40 years; >40) on of SARS-CoV-2 entrance/processing genes in nasal epithelial cells. Data is shown as individual points for each gene and analysed using Students T-Test or Mann-Whitney T-test as appropriated; significance was accepted when *p<0.05.

**Supplementary Figure 9:**
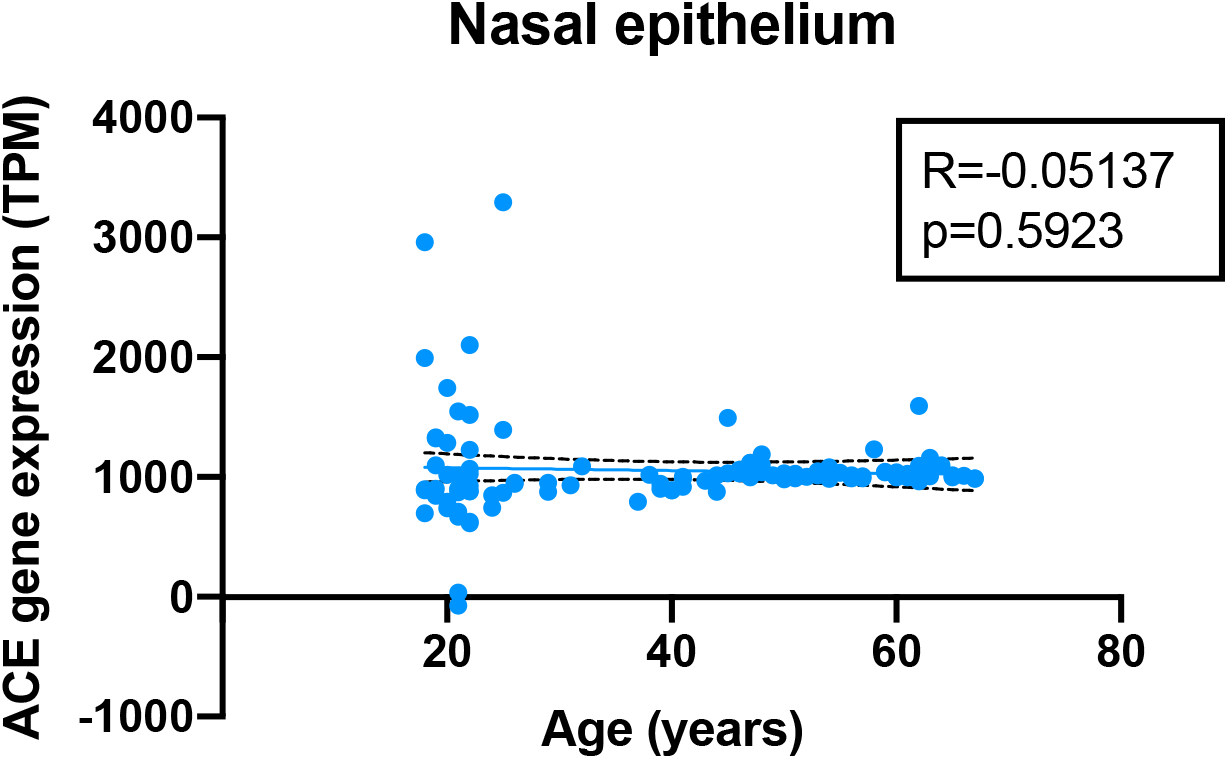
Correlation analysis of ACE expression in nasal epithelial cells with age. Data is shown as individual points for *ACE* expression levels and age (years). Data was analysed using Pearson’s correlation analysis.

**Supplementary Figure 10:**
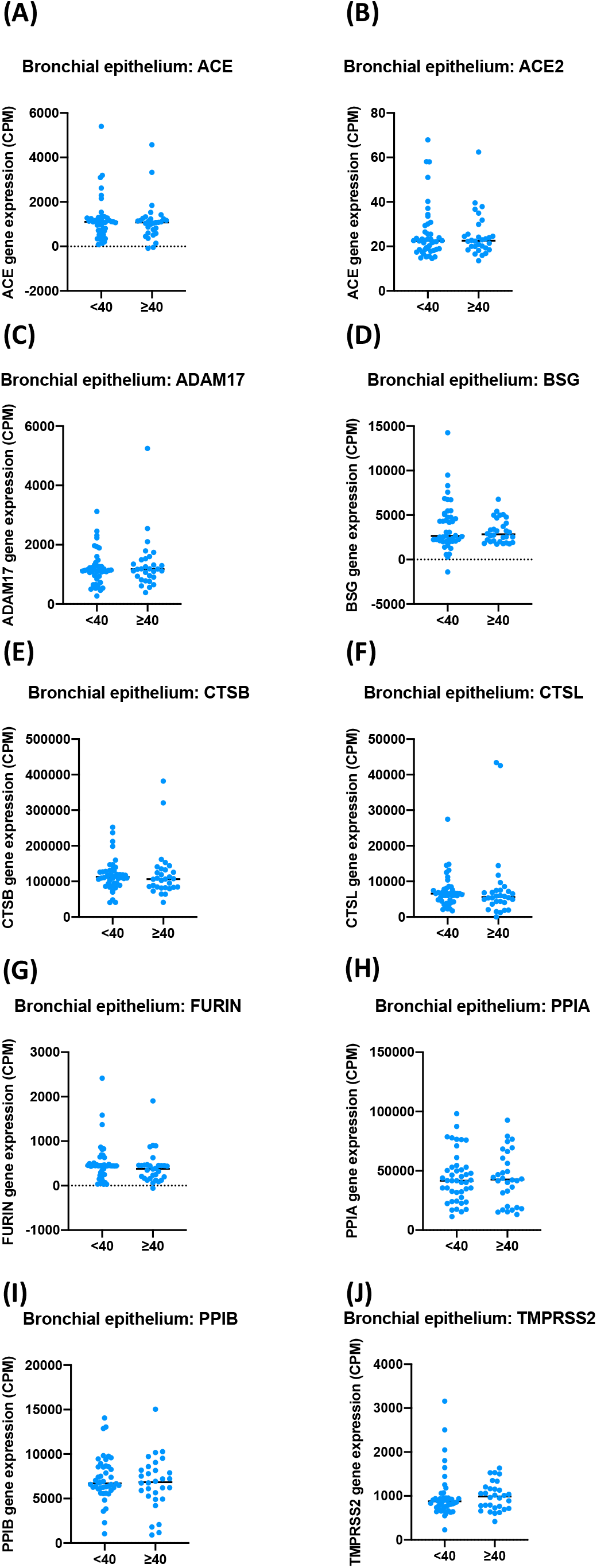
The effect of age (adult under 40 years; <40 versus over 40 years; >40) on of SARS-CoV-2 entrance/processing genes in bronchial epithelial cells. Data is shown as individual points for each gene and analysed using Students T-Test or Mann-Whitney T-test as appropriated; significance was accepted when *p<0.05.

**Supplementary Figure 11:** The effect of age (adult under 40 years; <40 versus over 40 years; >40) on of SARS-CoV-2 entrance/processing genes in peripheral blood mononuclear cells (PBMCs). Data is shown as individual points for each gene and analysed using Students T-Test or Mann-Whitney T-test as appropriated; significance was accepted when *p<0.05.

**Supplementary Figure 12:**
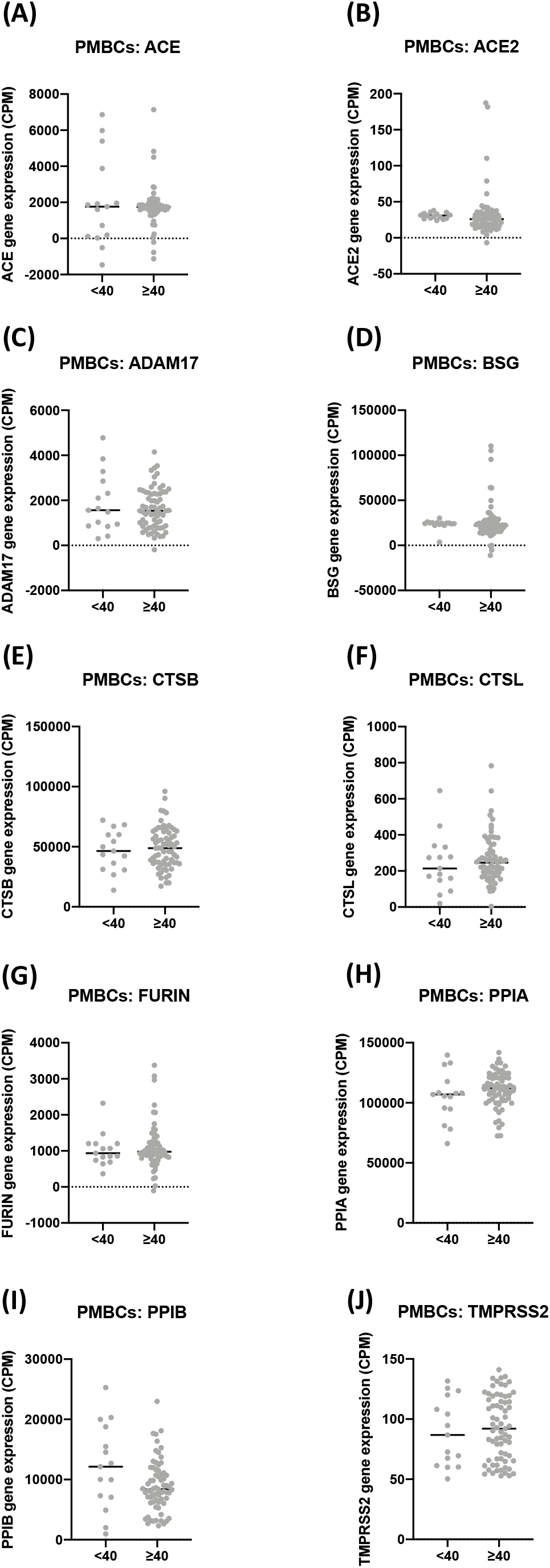

